# Reliable estimation of tree branch lengths using deep neural networks

**DOI:** 10.1101/2022.11.07.515518

**Authors:** Anton Suvorov, Daniel R. Schrider

## Abstract

A phylogenetic tree represents hypothesized evolutionary history for a set of taxa. Besides the branching patterns (i.e., tree topology), phylogenies contain information about the evolutionary distances (i.e. branch lengths) between all taxa in the tree, which include extant taxa (external nodes) and their last common ancestors (internal nodes). During phylogenetic tree inference, the branch lengths are typically co-estimated along with other phylogenetic parameters during tree topology space exploration. There are well-known regions of the branch length parameter space where accurate estimation of phylogenetic trees is especially difficult. Several novel studies have recently demonstrated that machine learning approaches have the potential to help solve phylogenetic problems with greater accuracy and computational efficiency. In this study, as a proof of concept, we sought to explore the possibility of machine learning models to predict branch lengths. To that end, we designed several deep learning frameworks to estimate branch lengths on fixed tree topologies from multiple sequence alignments or its representations. Our results show that deep learning methods can exhibit superior performance in some difficult regions of branch length parameter space. For example, in contrast to maximum likelihood inference, which is typically used for estimating branch lengths, deep learning methods are more efficient and accurate when inferring long branches that are associated with distantly related taxa and perform well in the aforementioned challenging regions of the parameter space. Together, our findings represent a next step toward accurate, fast, and reliable phylogenetic inference with machine learning approaches.

## Introduction

Phylogenetic inference is primarily concerned with the estimation of different evolutionary parameters from multiple sequence alignments (MSA) of molecular data using a wide range of statistical approaches. One of the major objectives is to estimate a tree topology, which can be viewed as unconventional parameter (Yang 2014) that determines historical relationships of the taxa under consideration. Other fundamental parameters of interest are the branch lengths that represent evolutionary (genetic) distances between the nodes of this tree. Typically, the branch lengths, ***b***, of a tree estimated from MSA’s represent relative distances expressed in terms of the time duration *t* and the substitution rate *θ*. The parameters *t* and *θ* are not identifiable individually (Dos Reis and Yang 2013; Rannala 2016) unless there is additional information available, such as fossil record, dated geological events and/or direct observation of mutation rates (Ho et al. 2011). Despite the wealth of research literature in phylogenetics that focuses on tree estimation methods, the problem of branch length inference traditionally receives less attention than topology inference. To estimate ***b*** under maximum-likelihood (ML) criteria, these branch lengths are simultaneously optimized on a fixed tree topology to achieve an optimal likelihood score (Felsenstein 1981). Typically, branch-length estimates are considered to be independent quantities, satisfying the Markov property of the substitution process under ML criteria (Lyons-Weiler and Takahashi 1999).

In recent years machine learning methods, and deep learning frameworks in particular, have gained an increasing amount of attention from the phylogenetic community. A wide range of phylogenetic tasks were tackled from the machine learning perspective including tree topology estimation (Suvorov et al. 2020; Zou et al. 2020), the investigation of zones of phylogenetic bias (Leuchtenberger et al. 2020; Suvorov et al. 2020), the identification of autocorrelation in evolutionary rates (Tao et al. 2019), model selection (Burgstaller-Muehlbacher et al. 2022), taxon placement on existing trees (Jiang et al. 2022) and improving tree topology search (Azouri et al. 2021).

Estimation of ***b*** is a critical step in phylogenetic inference because a great deal of statistical methods of divergence time estimation critically depend on the accuracy of these estimates (Schwartz and Mueller 2010) as absolute geological time *t* is confounded in ***b***. If ***b*** is not estimated accurately, then subsequent calibration analyses will in turn be incorrect and may result in erroneous inferences from a wide range of evolutionary models. These may include phylodynamic analyses of viral outbreaks where branch lengths provide critical information about lineage turnover (Attwood et al. 2022) or estimation of macroevolutionary parameters such as speciation and extinction rates from time-calibrated phylogenies (Marshall 2017). Thus, inference of ***b*** will exert a “domino effect”, whether positive of negative, on the numerous downstream evolutionary analyses.

Here, we provide a machine learning framework to estimate ***b*** ether directly from an MSA using convolutional neural networks, or from summaries of the MSA, such as site pattern frequencies, using a multilayer perceptron. We further compare the performance of these methods with the most commonly used method of branch length inference—the maximum likelihood approach. Our simulation study shows that machine learning methods provide reliable estimates of ***b*** often outperforming maximum likelihood and excelling in certain difficult regions of the branch-length parameter space.

## Methods

Our simulation procedures were divided into two consecutive tasks: (i) the simulation of tree branch lengths (***b***) and (ii) the generation of corresponding MSAs. Then, simulated ***b*** and MSAs were used to compare the performance of branch-length inference by maximum likelihood to our deep learning approaches. We examined a variety of different combinations of parametric distributions for ***b*** and models of the substitution process. We describe these in detail below.

### Uniform and exponential models

We simulated branch lengths for unrooted quartet topologies under several uniform distributions with different minimum and maximum parameter values: *U*(0, 0.001), *U*(0.001, 0.01), *U*(0.01, 0.1), *U*(0.1, 1) and *U*(1, 10). We also simulated ***b*** under an exponential distribution, which is viewed as a “biologically interpretable” probability model (Venditti et al. 2010). In particular, we examined three exponential distributions with mean (α) of 1, 0.1 and 0.01, respectively. Each branch length in a simulated tree was drawn independently from the same parametric distribution. Additionally, we used *U*(0.1, 1) and *U*(1, 2) distributions to simulate data for the “misspecified branch length distribution” experiments, whereas *Exp*(0.1) distribution was used to generate ***b*** for 8-taxon trees with different degrees of topology balance (**Table 1**).

**Table 1:** Summary of the multiple sequence alignment simulation experiments that were performed in this study using different branch length distributions, substitution models and tree topologies. *U*(*min, max*) = uniform distribution with the corresponding minimum and maximum parameter values. *Exp*(α) = exponential distribution with the corresponding mean parameter α. ϵ is the relative extinction parameter for the birth-death model (i.e. *μ*/*λ* where *μ* and *λ* are the death and birth parameters, respectively) and *a* is the age of the root node in Mya of the birth-death model. BL-space = branch-length heterogeneity space. Each simulation consists of a combination of a specific branch length distribution, substitution model and tree topology. For example, the “exponential” experiment includes a total of six parameter combinations: all pairs of the three branch length distributions and two substitution models in the “Exponential” entry in the table.

| Experiment | Branch length distribution parameters | Substitution models | Topology | Remarks |
| --- | --- | --- | --- | --- |
| Uniform | $U(0, 0.001)$, $U(0.001, 0.01)$,<br>$U(0.01, 0.1)$, $U(0.1, 1)$, $U(1, 10)$ | JC | unrooted 4 taxon<br>(A,B,(C,D)) | |
| Exponential | $Exp(1)$, $Exp(0.1)$, $Exp(0.01)$ | JC, GTR | unrooted 4 taxon<br>(A,B,(C,D)) | |
| BL-space | Beta mixture | JC, GTR | unrooted 4 taxon<br>(A,B,(C,D)) |  |
| Birth-death | $\epsilon = 0, 0.5$ or $0.9$<br>$a = 10, 50, 100$ or $200$<br>Clock rate = 0.001 | 3.3b,<br>UNREST | rooted 4 taxon<br>((A,B),(C,D)) | |
| Model misspecification | $Exp(1)$, $Exp(0.1)$, $Exp(0.01)$ | JC, GTR | unrooted 4 taxon<br>(A,B,(C,D)) | Train dataset was simulated under JC and applied to test dataset simulated under GTR and vice versa. |
| Misspecified branch length distribution | $U(0.1, 1)$, $U(1, 2)$ | JC | unrooted 4 taxon<br>(A,B,(C,D)) | Train dataset was simulated using $U(0.1, 1)$ and test dataset was simulated using $U(1, 2)$ and vice versa. |

| Effects of tree topology balance | Exp(0.1) | GTR | unrooted 8 taxon<br>(A,B,((C,D),((E,F),(G,H))))<br>(A,B,((C,D),(E,(F,(G,H))))))<br>(A,B,(C,(D,((E,F),(G,H))))))<br>(A,B,(C,(D,(E,(F,(G,H)))))) |
| --- | --- | --- | --- |

### Sampling from branch-length heterogeneity space

Additionally, to assess the impact of branch-length asymmetry on estimation accuracy, we simulated ***b*** from our previously described “branch-length heterogeneity space” (Suvorov et al. 2020) or “BL-space”. Briefly, to generate these data we simulated 10^6^ quartet topologies with randomly assigned branch lengths drawn from the following mixture of beta distributions:

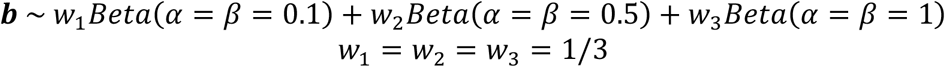

Then, we projected each tree branch length configuration onto a 2D space using three simple statistics: the sum of pairwise differences (PD) of branch lengths, total tree length (L) and sum of lengths of neighboring branches (NS) as detailed in Suvorov et al. (2020). Finally, we uniformly sampled branch length configurations from this 2D space. This approached allowed us to simulate trees with various degree of branch length heterogeneity, including trees from regions of the parameter space that are known to cause biased estimation of tree topologies, such as Felsenstein (Huelsenbeck and Hillis 1993) and Farris (Siddall 1998) zones.

### Birth-death model

We simulated trees under the birth-death (BD) process using a two-step procedure: 1) the generation of the absolute branch lengths ***b***_*a*_ using RevBayes v1.0.12 (Höhna et al. 2016), followed by 2) the conversion of ***b***_*a*_ to relative branch lengths ***b*** using a strict clock model in NELSI v0.21 (Ho et al. 2015). First, we generated ***b***_*a*_ from the following model:

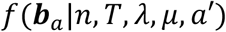

where *T* is a fixed tree topology, *n* is the number of taxa, *λ* and *μ* are the birth and death rates, respectively, and *a*′ is a root node age drawn from *U*(*a, a* + 0.001). This tight uniform prior was used because we desired a fixed root node age, but RevBayes requires a prior distribution. In order to constrain *T* in RevBayes, we disallowed all tree topology move proposals, leaving only the scale and slide operators that change branch lengths only. We did not specify *λ* and *μ* directly, but instead specified the relative extinction parameter, i.e. ϵ = *μ*/*λ*, and then computed *λ* and *μ* using the method-of-moments estimator (Magallon and Sanderson 2001) for the net diversification rate, *r*, as follows:

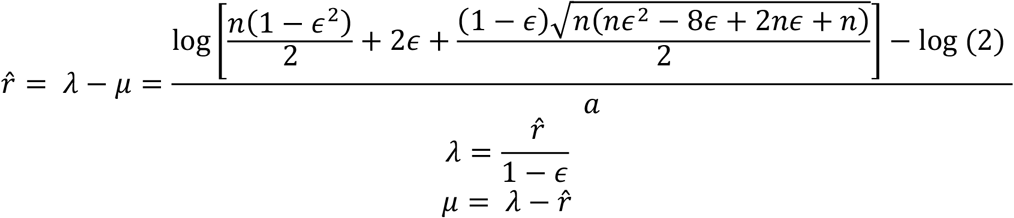

We used all pairwise combinations of the following BD parameter values: ϵ of 0, 05 and 0.9, and *a* of 10, 50, 100 and 200. Then, we performed Markov Chain Monte Carlo (MCMC) sampling in RevBayes to generate ***b***_*a*_ after discarding the first 5,000 generations as burn-in. Finally, ***b***_*a*_ were scaled to ***b*** in NELSI using strict clock model with a constant rate of 0.01 substitutions per site per million years, and no noise.

### Multiple sequence alignment simulations

MSAs were simulated using phylogenetic simulator AliSim (Ly-Trong et al. 2022) which is integrated into the IQ-TREE v2.2.0 software (Minh et al. 2020). We used a wide range of DNA substitution models including time-reversable Markov models such as JC and GTR as well as non-reversable Lie Markov models (Woodhams et al. 2015), namely 3.3b and UNREST (Yang 1994) (which is equivalent to 12.12 model in IQ-TREE). The free substitution rate parameters for GTR were drawn from *U*(0, 1), whereas for the UNREST model these parameters were drawn from *U*(0, 0.979) as they cannot exceed value of 0.98 for Lie Markov models (if this condition is not satisfied AliSim raises an error). The equilibrium base frequencies for JC and 3.3b were specified with 0 degrees of freedom, i.e. *π*_*A*_ = *π*_*G*_ = *π*_*C*_ = *π*_*T*_ = 1/4, whereas for GTR and UNREST base frequencies were unconstrained with 3 degrees of freedom and were drawn from *U*(0, 1) and then normalized in order to satisfy *π*_*A*_ + *π*_*G*_ + *π*_*C*_ + *π*_*T*_ = 1. The among-site rate variation (+Γ) was used with every substitution model and was modeled using discrete Gamma model with four rate categories with the shape parameter drawn from *U*(0, 1). Thus, every MSA was simulated along a given tree (with branch lengths drawn as described above) using the specified substitution model parameters. Our complete set of simulation experiments and their respective combinations of branch length distributions, substitution models and tree topologies are summarized in **Table 1**.

### Deep learning architectures and training

For each simulation experiment shown in **Table 1**, we generated a training set consisting of 15 × 10^4^ trees with branch lengths drawn from a specified distribution, with the MSA for each tree then generated using AliSim. The length of each MSA was set to 1,000 sites. We treated each MSA as a matrix with rows corresponding to sequences and columns corresponding to sites in the alignment, and with each character in the alignment encoded as a number (“A”:0, “T”:1, “C”:2, “G”:3).

The main objective of a supervised machine learning algorithm is to learn mapping of inputs, ***X***, to outputs, ***Y***. In our case the MSAs, or a representation thereof, served as inputs, and the branch lengths ***b*** were our outputs. Since the goal is to predict positive real-valued ***b*** ∈ ℝ ^+^ from a given MSA, our task is a regression problem. Within this study we explored and compared behavior of two types of artificial neural networks trained to complete this task: a multilayer perceptron (MLP) and a convolutional neural network (CNN). We used the Keras v2.9.0 Python API (https://keras.io/) with TensorFlow v2.9.1 as the backend (Abadi et al. 2016) to build, train and test these machine learning models.

As our input for the MLP, we constructed feature vectors ***X*** by counting site patterns found within an MSA and then divided these counts by the alignment length. There exist exactly 4^4^ = 256 such possible patterns (i.e., features) for an alignment of four taxa, as gap characters were not examined in this study. For the “effects of tree topology balance” experiments we extracted site patterns directly from 8-taxon alignments resulting in 4^8^ = 65,536 features. Our MLP architecture comprised an input layer, three fully connected dense hidden layers of 1,000 neurons each with rectified linear unit (ReLU) activation, and an output layer with the number of neurons set to the number of tree branches and with linear activation. Additionally, we used the dropout regularization technique with a rate of 0.15 applied to every hidden layer.

The CNN architectures built for this study used network architectures/hyperparameters similar to the ones utilized in (Suvorov et al. 2020) with some modifications. The CNN uses entire MSAs as input ***X*** and extracts abstract features by performing several convolutional and pooling operations. Our CNN contained an input layer, six convolutional layers consisting of 1024, 1024, 1024, 128, 128 and 128 filters, respectively. Each convolutional layer used ReLU activation and was followed by a pooling layer. We set the kernel size for the first convolution operation to 4 × 1, reasoning that it would potentially capture a phylogenetic signal by striding across the MSA and examining one column of the alignment at a time, and the subsequent convolution steps had each had kernel sizes of 1 × 2. The first average-pooling operation size was set to 1 × 1 (i.e., no pooling), with subsequent pooling steps having sizes of 1 × 4, 1 × 4, 1 × 4, 1 × 2 and 1 × 2. Finally, the output of the last pooling layer was flattened and passed to a fully connected dense layer of 1,000 neurons with ReLU activation, which in turn was connected to the output layer, whose number of neurons was equal to the number of tree branches. Again, the output layer used linear activation. Here, we did not use batch normalization or dropout regularization, as we observed that these tended to negatively affect learning stability and the accuracy of predicted branch lengths.

Machine learning regression models can suffer from systematic biases (Belitz and Stackelberg 2021; Igel and Oehmcke 2022) caused by overestimation and underestimation of small and large output variables ***Y***, respectively, and/or by a transformation of original output variables ***Y***. We observed the latter form of bias in our initial experiments, and therefore sought to correct this using the “regression of observed on estimated values” (ROE) technique (Belitz and Stackelberg 2021). This approach includes two steps: (i) a *trained* neural network model is used to predict output values ***Ŷ*** on training dataset ***X***, and (ii) these estimated values ***Ŷ*** from the training set are used in turn to train a separate regression model seeking to better estimate the true output values ***Y***. To build and train this second regression model, we used a simple linear regression model implemented in Keras (i.e. an input layer, whose size is equal to the number of tree branches, connected to an output layer of the same size and which uses linear activation).

When training our MLP, CNN, ROE regression models, 10% of the training dataset was set aside for validation. We used mean squared error (MSE) as our loss function. We initially found that our neural networks occasionally predicted negative branch lengths whose values were less than zero. We therefore we performed square root transformation of ***b*** prior to training, and then squared the predicted ***b*** after running the network, thereby ensuring that predicted ***b*** ∈ ℝ^)^. The networks’ parameters (weights) were updated during training using the adaptive moment estimation (Adam) optimizer with batches sizes of 100, 32, and 32 for the MLP, CNN and ROE architectures, respectively. To prevent a neural network from overfitting, we used early stopping by monitoring validation loss metric during training and used a patience value of 10, causing training to terminate whenever loss on the validation set failed to improve for 10 consecutive training epochs. Loss was considered to have improved if it decreased by at least 0.0001 relative to the minimum loss observed across all previous epochs in the run (i.e. min_delta=0.0001).

In total, we constructed four deep learning models: MLP, CNN, MLP-ROE, and CNN-ROE where the latter two were trained by adding the linear regression step onto the previously trained MLP and CNN regression models. The flowchart in **Fig. 1** summarizes the steps of the training procedure and the nature of our input and target variables.

**Figure 1:**
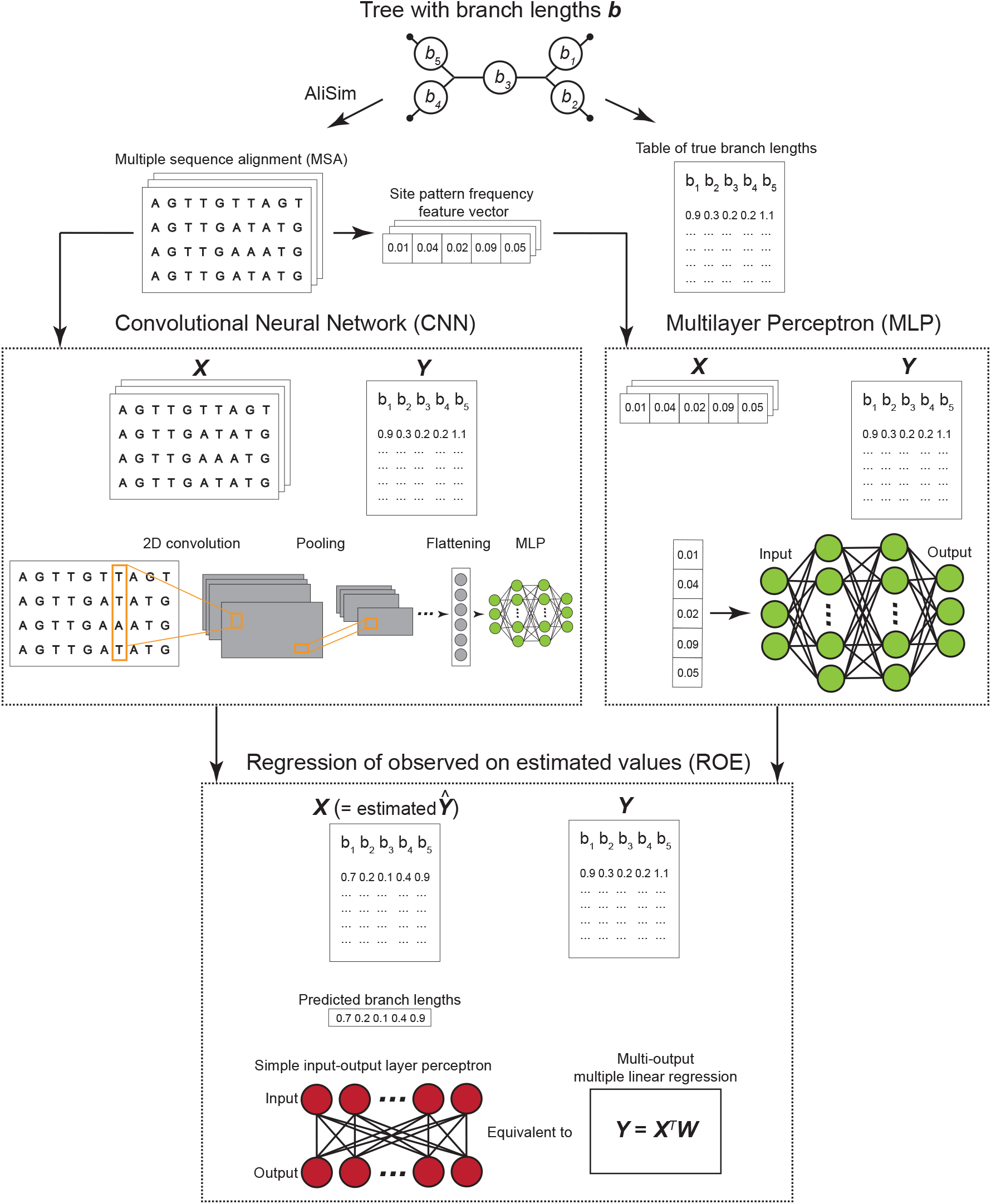
Flowchart summary of simulation and artificial neural network training procedures. After generating trees with branch lengths, multiple sequence alignments (MSA) were simulated with AliSim. From each MSA site pattern frequencies were calculated. MSAs (or site frequency vectors) together with the true branch lengths were used as input ***X*** and target ***Y*** in the convolutional neural network (CNN) and multilayer perceptron (MLP) architectures. The outputs Ŷ estimated by the CNN (or MLP) on the training dataset were used as an input ***X*** in the additional “Regression of observed on estimated values” step, where the true values were again used as target ***Y***. Once all networks were trained, they were subsequently used to predict branch lengths from a testing dataset.

### Testing procedures

We compared the performance of the deep learning architectures described above to a standard method of branch length tree estimation: maximum likelihood (ML) estimation as implemented in IQ-TREE. To that end, for each simulation experiment we generated an additional test set of 10,000 trees, each with branch lengths ***b***, and their corresponding MSA’s. In order to estimate branch lengths on a fixed tree topology in IQ-TREE, we generally used the same substitution model that had been specified to generate the input MSA. In the “Model misspecification” experiments (see **Table 1**), however, we used a different substitution model in IQ-TREE, and for training our networks, than that of the test data, allowing us to observe the impact of model misspecification on branch-length estimation accuracy. Again, we also used branch length distributions that differ between the training and test data for the “Misspecified branch length distribution” experiments, although we note that this has no impact on maximum likelihood estimation, which does not require a pre-specified distribution of branch lengths.

We summarized the performance of each method by measuring the mean squared error (MSE) and mean absolute error (MAE) for each estimated branch length and total tree length (i.e., the sum of the branch length vector ***b***). MSE is a useful criterion to compare efficiency of estimators, i.e. our branch length inference methods, where lower MSE values indicate higher efficiency. Additionally, we quantified bias by comparing the empirical cumulative distribution functions (eCDF) of observed and estimated ***b***. First, error (*ε*) was calculated using the sample quantile function *Q* for 100 quantiles of the true branch lengths:

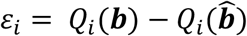

Where ***b*** and 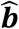 represent the true and estimated branch lengths, respectively, and *i* denotes the *i*^th^ quantile. Then, bias was computed as:

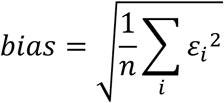

where *n* is equal to the total number of quantiles (i.e. 100). Further, the distributions of observed and estimated ***b*** were statistically compared using a two-sample Kolmogorov-Smirnov (KS) test. We report KS test statistic *D*, which together with *bias* measure (largest) vertical and (average) horizontal distances, respectively, between true and observed eCDF’s. The relationships between observed and predicted values of ***b*** were also evaluated using the Spearman rank correlation test (SR) and its estimated correlation coefficient *ρ*. We also examined whether our methods tend to overestimate or underestimate ***b*** within certain parameter spaces. Specifically, we compared distributions of residuals, i.e. differences between true and estimated ***b***, by calculating the ratio of residuals with “–” and “+” signs, where the ratios of >1would suggest overestimation, whereas the ratios of <1 are indicative of underestimation. The significance of this ratio was assessed using a sign test.

## Results and Discussion

### ANNs accurately infer branch lengths generated by uniform distributions

We sought to compare the performance of several artificial neural network (ANN) estimators of branch lengths to that of maximum likelihood (ML) estimation (see Methods). First, as a simple scenario we evaluated each methods’ performance on the tasks of estimating the branch lengths ***b*** of a quartet tree sampled from generic uniform distributions with different minimum and maximum parameter values and MSAs simulated under JC model (see **Table 1**). In these experiments, the branch lengths ranged from very short (averaging 0.0005 substitutions per site) to very long branches (averaging 5.5 substitutions per site). Biologically, such contrasting scenarios reflect extremely slow versus fast uniformly evolving sequences or alternatively they can represent recent radiations versus deep taxonomic divergences. From a methodological perspective these scenarios encompass difficult parameter spaces, because MSAs simulated along very short trees will have limited phylogenetic information, whereas MSA’s simulated using long trees will be highly saturated with substitution events, which will erode phylogenetic signal in sequences that constitute the MSA, making them all appear highly dissimilar (Lee 2001). Thus, both of these scenarios may compromise a method’s ability to infer tree topology and ***b*** accurately (Yang 1998; Duchêne et al. 2022). For example, it is expected that multiple hits, if not accounted for, (Yang et al. 1994; Schwartz and Mueller 2010) or cases of “deep-time” substitutional saturation (Moody et al. 2022) will lead to underestimation of ***b***.

For the *U*(0, 0.001) experiment, ANNs outperformed ML across all branches and total tree lengths (supplementary table S1). Interestingly, the ANNs tended to infer branch lengths closer to the mean of *U*(0, 0.001) which is equal to 0.0005 (supplementary fig. S1a). ML, on the other hand, showed exceptionally poor performance with highly inaccurate branch predictions (supplementary table S1). All methods underestimated branch lengths (supplementary fig. S2a). For the *U*(1, 10) experiment we observed that all ANNs exhibit similar performance across branches and tree lengths (**Fig. 2a**; supplementary table S1), whereas ML shows markedly inferior inferences (supplementary table S1). For example, for total tree length estimates the MSE, MAE, Spearman’s *ρ*, and *bias* for all ANN architectures (with the marginal superiority of MLP and MLP-ROE) were ~28, ~4.2, ~0.44 and ~3.6, respectively but ML’s measures were ~154, ~10, ~0.27 and 9.2, respectively. In this case, as a general trend we noticed that all methods underestimate ***b*** (**Fig. 2b**), however for branches 2 and 3 CNN-ROE did not show any such systematic bias. A comparison of the true and estimated means of branch length distributions reveals that the average ***b*** and total tree length was inferred relatively accurately by ANNs (**Fig. 2c**). ML, on the other hand, underestimates ***b***, so that the estimated density of ***b*** is mostly concentrated near the lower boundary of *U*(1,10) with some predictions being even lower (**Fig. 2c**). In other experimental scenarios (supplementary table S1; supplementary data), the ANNs perform well, and almost always outperformed ML or exhibited similar accuracy.

**Figure 2:**
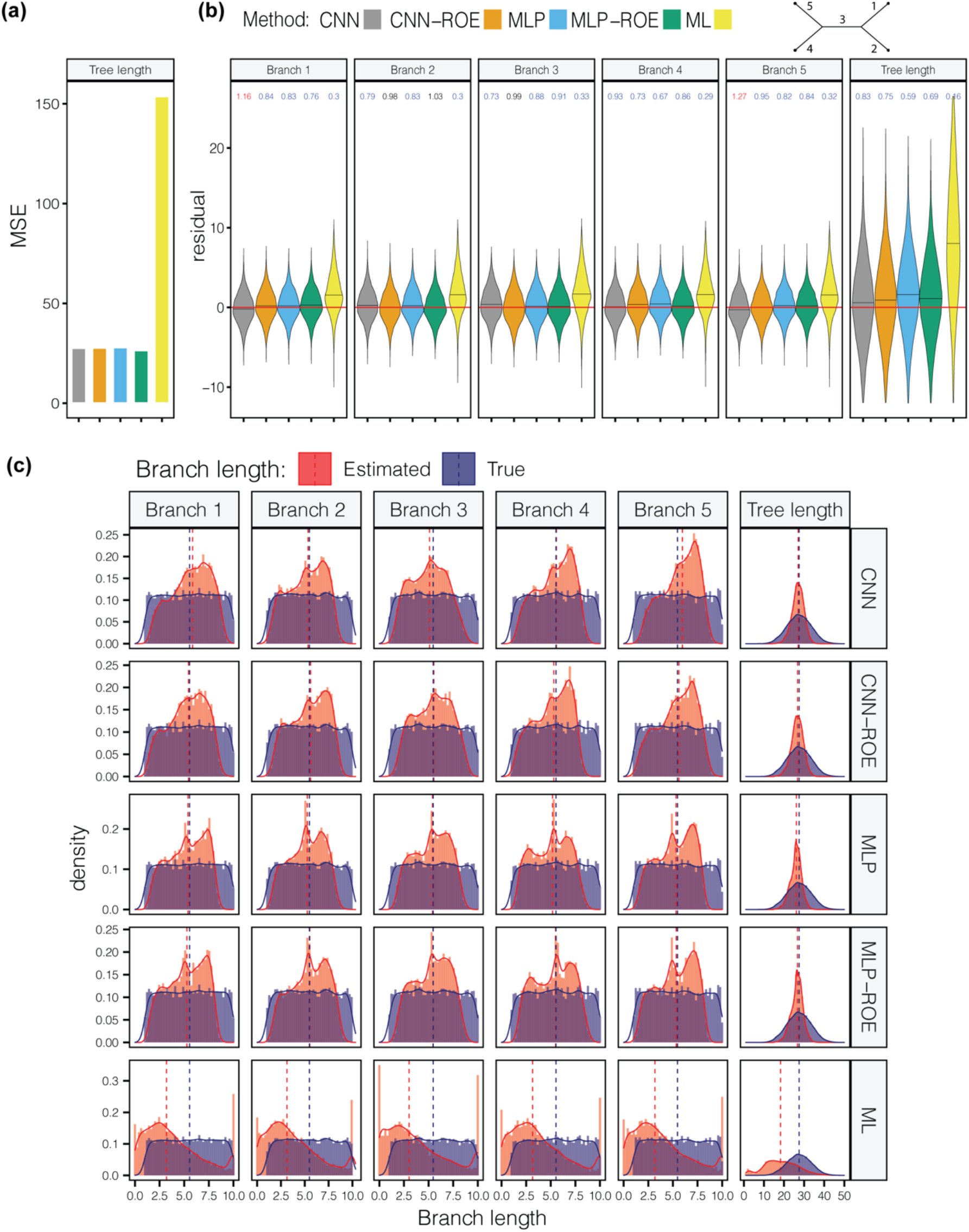
Performance of methods for long branch lengths generated using uniform distribution with minimum and maximum parameters of 1 and 10, respectively. (**a**) Comparison of mean squared error (MSE) for all methods. All artificial neural network models show superior performance relative to maximum likelihood. (**b**) The violin plot that shows the distribution of residuals (i.e. difference between true and inferred branch lengths) for each method. The value above each violin represents the ratio of overestimated and underestimated branches. The values of ~1 indicate an equal number of over- and underestimates, <1 indicate the underestimation is more common, and >1 indicates that overestimation is more common. The colored values represent statistically significant underestimation (blue) or overestimation (red). The horizontal red line marks 0. The black horizontal line within each violin shows the median. (**c**) Comparison of true (blue) and predicted (red) branch length distributions. Dashed lines mark the position of the median. CNN = convolutional neural network; CNN-ROE = convolutional neural network – regression of observed on estimated values; MLP = multilayer perceptron; MLP-ROE = multilayer perceptron – regression of observed on estimated values; ML = maximum likelihood.

Together, the distribution of residuals and comparison between true and predicted ***b*** distributions can inform about the tendency and magnitude of a method’s over- or underestimation of ***b***. As a general trend, we noticed that ML produces an excess of severely overestimated ***b***, which can be seen in the left tail of predicted ***b*** distribution that goes beyond the boundaries (across all simulation scenarios that use uniform distribution) of the true ***b*** distribution (supplementary fig. S1 a-e). Interestingly, that ANNs tend to generate predicted ***b*** distributions within the theoretical distribution boundaries, which suggests that ANNs are not inclined to severely overestimate/underestimate ***b***. However, our examination of residual plots (supplementary fig. S2 a-e) revealed that in most of the cases, all methods are prone to underestimating ***b*** although we stress that the magnitude of underestimation is substantially lower for the ANNs than ML. Additionally, ANNs are more efficient estimators of ***b***, since without any exception across all the experiments with uniform distributions ANNs have always smaller MSE values for all branch lengths and the total tree length (supplementary table S1).

### ANNs accurately infer branch lengths generated by exponential distributions

We next evaluated each method’s performance on trees whose branch lengths were drawn from exponential distributions with means of 0.01, 0.1 and 1. The exponential distribution is commonly used as a prior for ***b*** in phylogenetic Bayesian frameworks (Ronquist and Huelsenbeck 2003) and is more biologically meaningful (Brown et al. 2010; Venditti et al. 2010) than a uniform distribution. For each of these exponential branch length distributions, we generated two sets of MSAs: one using the JC substitution model and one using the GTR model (see **Table 1**). Generally, methods’ performances are similar between experiments that use JC and GTR models, but with slightly better outcomes achieved for the former case (supplementary table S2; supplementary data). Here we describe the results obtained for the MSAs generated under JC. The performance across ANNs and ML was comparable for the experiment with exponential means of 0.01 and 0.1 (supplementary table S2). However, for the experiment with exponential mean of 1, ANNs significantly outperformed ML, showing error rates as measured by MSE that were an order of magnitude lower than that of ML (**Fig. 3**). The disparity between MAEs was smaller, which is best explained by ML’s tendency to on occasionally produce wildly inaccurate predictions, which have a larger impact on MSE than MAE. Interestingly, for all methods except the CNN without ROE (see Methods), the eCDF of predicted total tree lengths matched the true eCDF of total tree lengths (KS test, all *P* > 0.05); however, this result varied for individual branch lengths (supplementary table S2; supplementary data).

**Figure 3:**
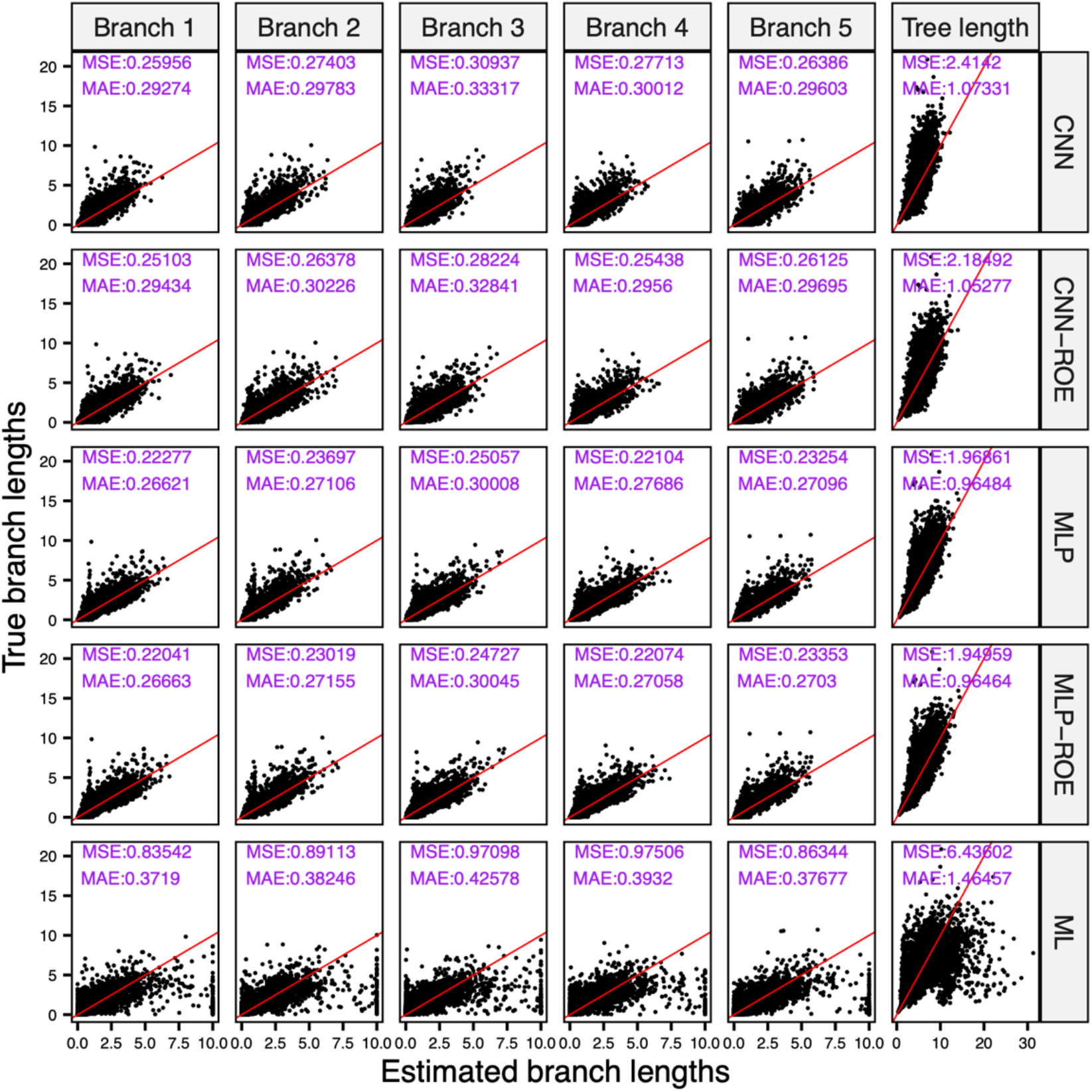
Correlation between estimated and true branch lengths drawn from exponential distribution with mean of 1. All artificial neural network models show higher efficiency and accuracy than the maximum likelihood method. CNN = convolutional neural network; CNN-ROE = convolutional neural network – regression of observed on estimated values; MLP = multilayer perceptron; MLP-ROE = multilayer perceptron – regression of observed on estimated values; ML = maximum likelihood; The numbers in purple indicate mean squared error (MSE) and mean absolute error (MAE). Red line indicates perfect linear relationship.

### ANNs show elevated accuracy for trees with long branches

Trees with heterogeneous branch lengths can significantly deteriorate the performance of phylogenetic methods. Classical examples of such regions of branch length parameter space include the Felsenstein zone (Huelsenbeck and Hillis 1993) and the Farris zone (Siddall 1998), where even statistically consistent methods may not exhibit good accuracy in tree topology estimation, depending on an MSAs length. In fact, deep learning approaches may also suffer in such regions, especially in the Felsenstein zone (Suvorov et al. 2020). However, this drawback can be ameliorated by increasing the alignment length, and in the case of deep learning methods, altering the composition of the training set to include more examples from these challenging regions of the parameter space. Inference of branch lengths can be also affected within these regions: it has been shown that branch length heterogeneity can decrease a method’s efficiency (i.e. its accuracy on smaller MSAs) and cause estimation artifacts such as non-independence of inferred ***b*** (Lyons-Weiler and Takahashi 1999).

We previously described a distribution of branch lengths that captures a wide array of tree configurations, including those in the Farris and Felsenstein zones and other challenging regions of the parameter space. This distribution, which we call the BL-space (**Fig. 4a**), is described in detail in Suvorov et al. (2020) and briefly in the Methods section. Our simulations show that ANNs exhibit overall significantly better performance on MSAs with branch lengths drawn from this BL-space in both JC and GTR testing datasets in terms of the MSE, MAE, *ρ* and *bias* across all branches of our quartet topologies and the total tree lengths (supplementary table S3; supplementary data). Although none of the examined methods were able to produce distribution of ***b*** estimates that match true distribution of ***b*** in BL-space (KS test, all *P* < 0.05), the *bias* and *D* statistic were noticeably smaller for ANNs than for ML (supplementary table S3). We also note that ROE consistently improves the accuracy of MLP and CNN as measured by MSE, MAE and *bias* (supplementary table S3). More detailed comparison of the true and estimated ***b*** distributions revealed that ANN models tend to infer ***b*** that more closely follow the shape of the true distribution of tree lengths (**Fig. 4b**) as well as the shapes of ***b*** distributions in individual branches (supplementary fig. S3), than do the estimates from ML. Moreover, ***b*** predicted by ANNs do not significantly extend beyond the domain of the BL-space, i.e. ***b*** ∈ [0,1] and tree lengths ∈ [0,5], whereas noticeable fraction of ML estimates goes beyond the upper bound of the BL-space domain. This observation suggests ***b*** overestimation by ML, but also that ANNs do not frequently produce estimates outside of their training range, a notion we examine further below. Then, we investigated the distribution of relative error 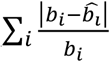, where *b* and 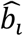 are true and estimated values for branch *i* for each tree) to see if methods show some performance differences affected by branch length heterogeneity that is not governed merely by the size of the branch. Consistent with our previous results for the uniform distributions, ANNs tend to produce lower relative estimation errors than ML for trees with longer branches (**Fig. 4c**). This observation is exemplified by the panels which show the density of trees that had the lowest relative estimation error (error < 25% quantile), which is highest towards the top of the BL-space for ANNs, where trees consisting entirely of long branches reside (**Fig. 4c**). Although, we did not observe any dramatic performance biases, all estimation methods had larger errors (error > 75% quantile) for the trees with small ***b*** on average and within the Felsenstein zone (**Fig. 4c**, bottom and lower middle parts, respectively, of the BL-space). These results are not surprising since trees with very short branches used in our simulations are expected to generate MSAs with very few variable sites, whereas Felsenstein zone is known to cause long branch attraction artifact for topology estimation problems and may also result in inaccurate ***b*** inference.

**Figure 4:**
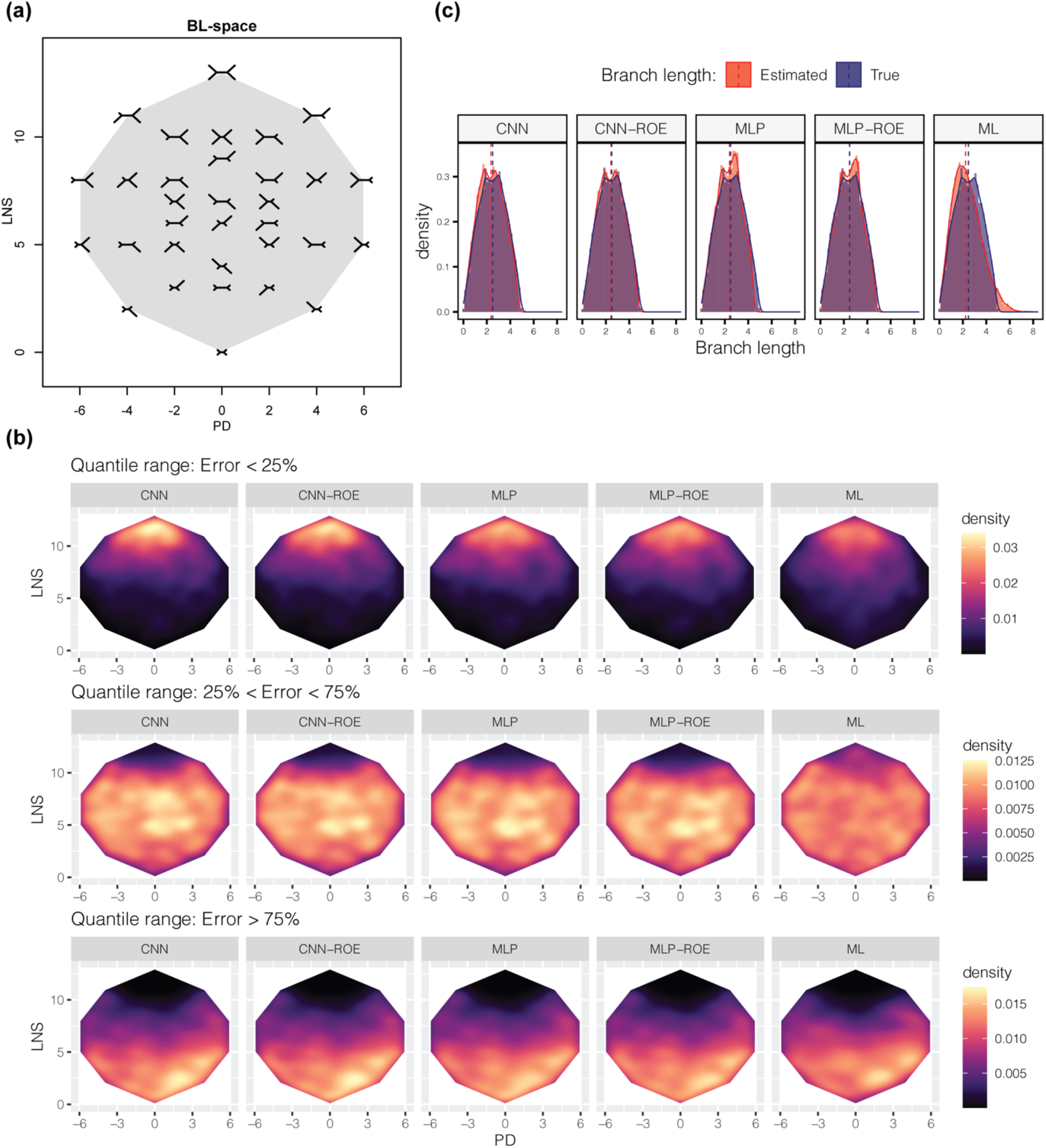
Performance within branch length heterogeneity space (BL-space). (**a**) The branch length space (BL-space). Each tree in the BL-space marks the location of the extreme cases where short branches in the image represent branch lengths equal to 0 and long branches represent lengths of 1. The shaded area shows the boundaries of the BL-space. PD = pairwise difference, LNS = tree length (L) + lengths of neighboring branches (NS). See Suvorov et al. (2020) for more detail. (**b**) Comparison of true (blue) and predicted (red) total tree length distributions. Dashed lines mark the position of the median. (**c**) Distribution of average tree errors (absolute value of a difference between true and predicted branch lengths divided by the true branch length). Each panel corresponds to error quantile ranges. CNN = convolutional neural network; CNN-ROE = convolutional neural network – regression of observed on estimated values; MLP = multilayer perceptron; MLP-ROE = multilayer perceptron – regression of observed on estimated values; ML = maximum likelihood.

### ANNs accurately estimate branches that connect to a root node

We simulated branch lengths using birth-death models assuming a strict clock on a rooted quartet topology with a balanced tree shape i.e. ((A,B),(C,D)). Since this process generated rooted trees, the MSAs were generated using 3.3b and UNREST substitution models which are non-reversable and thus are able to account for the position of a root node that splits a branch into two branches with identifiable lengths (Bettisworth and Stamatakis 2021).

Since the overall performance of ANNs and ML on MSAs generated under the 3.3b or UNREST models were similar (supplementary table S4; supplementary data), here we report the results only for model 3.3b. Additionally, the relative extinction parameter ϵ did not have a large effect on these methods’ performance (supplementary table S4). ANNs exhibited superior estimation accuracy compared to ML. Specifically, ML struggled to infer the lengths of branches that connect to the tree root node under non-reversible substitution models. The predicted ***b*** that are connected to a root node by ML were biased and had notably poorer correlation (however significant, SR test, all *P* < 0.05) with true root node ***b***, as exemplified by coefficients *ρ*, than those estimated by ANNs (supplementary table S4). The ML estimates of terminal branches were notably better than internal ones across all parameter settings but still inferior to the ANNs’ estimates. In contrast, ANNs were able to accurately infer all ***b***, including the lengths of internal branches connected to the root, which in turn implies that ANNs are capable of inferring the position of the root of a tree. The correlations between predicted and true ***b*** values, as well as between predicted and true total tree lengths, were strong and significant (SR test, all *P* < 0.05) with *ρ* > 0.92 across all simulation scenarios. In most of the cases the MSE and MAE metrics for ANNs were an order of magnitude lower than those reported for ML (supplementary table S4). All methods of branch-length estimation exhibited better performance for trees with root node age of 50 Mya or 100 Mya (supplementary table S4). This result is expected, because for these trees the average length of the path from the terminal node to the root node will be 0.05 substitutions/site or 0.1 substitutions/site, respectively. Under such parameter settings, simulated MSAs will have a sufficient number of variable sites but will not be too diverged to reliably estimate ***b***, as opposed to the other cases we tested (root ages of 10 Mya and 200 Mya) which have average path lengths of 0.01 or 0.2, respectively, which may produce less informative MSAs. Also, we note ANNs correctly infer that sister terminal branches (i.e. A and B or C and D) should have identical lengths as a result of tree ultrametricity resulting from our simulations that use strict clock. Specifically, our analysis of ***b*** predicted for A and B (or C and D) in the scenario with ϵ = 0.5, *a* = 100 show that their lengths are highly correlated (SR test, *ρ* ∼ 1, *P* = 0).

The notion that ANNs are able to precisely infer the tree root is particularly encouraging, since the problem of finding an appropriate outgroup remains a challenging task in practice (Pearson et al. 2013; Dang et al. 2022). Moreover, ANNs recognize clock-like ***b*** directly from MSAs, a property that can be potentially used to identify clock models directly from MSAs without the need to reconstruct phylogenetic trees. In fact, recent study shows that a machine learning framework can effectively distinguish between independent and autocorrelated branch rate models using summary statistics extracted from the trees (Tao et al. 2019). An advantage of using CNN architectures for this task is that they would not require any pre-defined summary statistics.

### Model misspecification and performance of ANNs outside of training parameter space

Here we describe evaluation of our methods under two conditions of model/parameter misspecification: (i) misspecification where training MSAs were generated under JC, whereas testing MSAs were generated under GTR (and vice versa) and (ii) testing ANNs on scenarios where the distribution of branch lengths used during training differs from that of the test set (**Table 1**).

Model selection plays an important part in phylogenetic inference. If a chosen model is a poor fit to molecular data, it may lead to erroneous inference of tree topology and/or branch lengths, (but see Spielman 2020). Although, topology estimation is less sensitive to model misspecification if the most parameter-rich model is specified, branch-length estimation can be more sensitive (Abadi et al. 2019). In general, all methods’ performance was less impacted by model misspecification in case when training MSAs were generated under GTR and testing MSAs were generated under JC than in the opposite scenario (supplementary table S5). This result is not surprising since the JC model is nested within the GTR model. Interestingly, we found that the CNN and CNN-ROE approaches were more susceptible to model misspecification than both ML, MLP and MLP-ROE methods, especially with ***b***~*Exp*(0.01) (supplementary table S5; supplementary data). A similar trend was observed for ***b*** inferred by ML. In the opposite setting, when MSAs simulated under GTR models but tested on MSAs generated under JC, however, ML produced comparable results with the ANNs. ANNs also exhibited greater robustness to model misspecification and produced notably better estimates than those produced by ML for the other two branch length distributions, i.e. *Exp*(1) and *Exp*(0.1).

Next, we examined the impact of misspecification of the distribution of branch lengths used during training. The ANNs showed unsatisfactory behavior in experiments where we trained ANNs on MSAs simulated using trees with ***b*** sampled from *U*(0.1, 1) and tested on *U*(1, 2) and vice versa. In both cases, the ANNs failed to accurately branch lengths, with the distributions of estimated ***b*** falling entirely outside of the true ***b*** distribution with no overlap (supplementary fig. S4; supplementary table S6). Thus, when using ANNs one must take care to ensure that the distribution of branch lengths used in training is not too narrow, so that it is likely to encompass the true ***b***.

### Effects of tree-shape balance on branch-length estimation

Besides branch-length heterogeneity, the degree of tree-shape balance, i.e. the branching pattern, may pose problems for inference of branch lengths, especially for those branches that are situated deeper in a tree (Colijn and Plazzotta 2018). To investigate this, we first evaluated the performance of each method across all 8-taxon tree shapes, branches as well as total tree lengths using MSE and MAE (**Fig. 5**). In general, for all examined 8-taxon tree shapes (**Table 1**) ANNs produced at least 4-fold lower MSE values than ML. ML, on the other hand, had slightly lower MAE values (at most 1.32-fold lower) than the ANNs across all branches (**Fig. 5**; supplementary table S7; supplementary data). This observation suggests that although ML typically infers branch lengths slightly more accurately than ANNs, ML occasionally grossly mis-infers a branch’s length, which is results in large MSE values. We note, however, that for quartet trees with branches drawn from *Exp*(0.1) and corresponding MSAs generated under GTR model (i.e. the same simulation conditions that were used for 8-taxon trees), the CNN-ROE, MLP and MLP-ROE outperformed ML both in terms of MSE and MAE metrics. For instance, total tree length MSE and MAE from MLP-ROE are ~9.07 and ~1.16 times better, respectively, than those obtained by ML (supplementary table S2; supplementary data). Thus, the marginally better values of MAE for ML are observed only for our 8-taxon trees and not the smaller 4-taxon trees. Also, while comparing performance of different ANNs on 8-taxon trees, we noticed that MLP-based architectures tend to be marginally more accurate than CNNs (**Fig. 5**).

Next, for each method we examined the distribution of MSE and MAE values and their variances across all branches to see whether the ***b*** estimates show any variation in accuracy depending on the degree of tree balance and branch depths (**Fig. 6**; supplementary table S8). As a general trend, we observed that MSE variance was nearly three orders of magnitude smaller for all ANNs than for ML, however the variance in MAE was similar or slightly smaller for ML when compared to ANNs, with the exception of MLP-ROE (supplementary table S8). Specifically, the MLP-ROE approach showed at least 1.2 times smaller MAE variance for all tree shapes compared to ML. Overall, we did not observe a systematic impact of tree shape on ***b***-estimation in terms of variance in MSE (or MAE): as we move from more symmetric to more asymmetric tree shapes in **Fig. 6**, we do not observe an increase in error. Nevertheless, we observed a strong tendency for internal branches in a tree to have higher errors for ***b***-estimates in comparison with terminal branches for both the ANNs and ML. This pattern was especially clear in the distribution of MAE across tree branches (**Fig. 6**). Taken together, our analyses suggest that 8-taxon trees represent a more difficult inferential problem than 4-taxon trees for ANNs, but on the other hand, ANNs on average showed adequate performance, and their superior MSE implies that they are highly reliable estimators in the sense that they do not produce the large errors sometimes observed for ML. We note here that it may be possible to further improve the ANN’s accuracy by constructing more optimal network architectures and/or using larger amounts of training examples simulated from the same parameter space (Suvorov et al. 2020).

**Figure 5:**
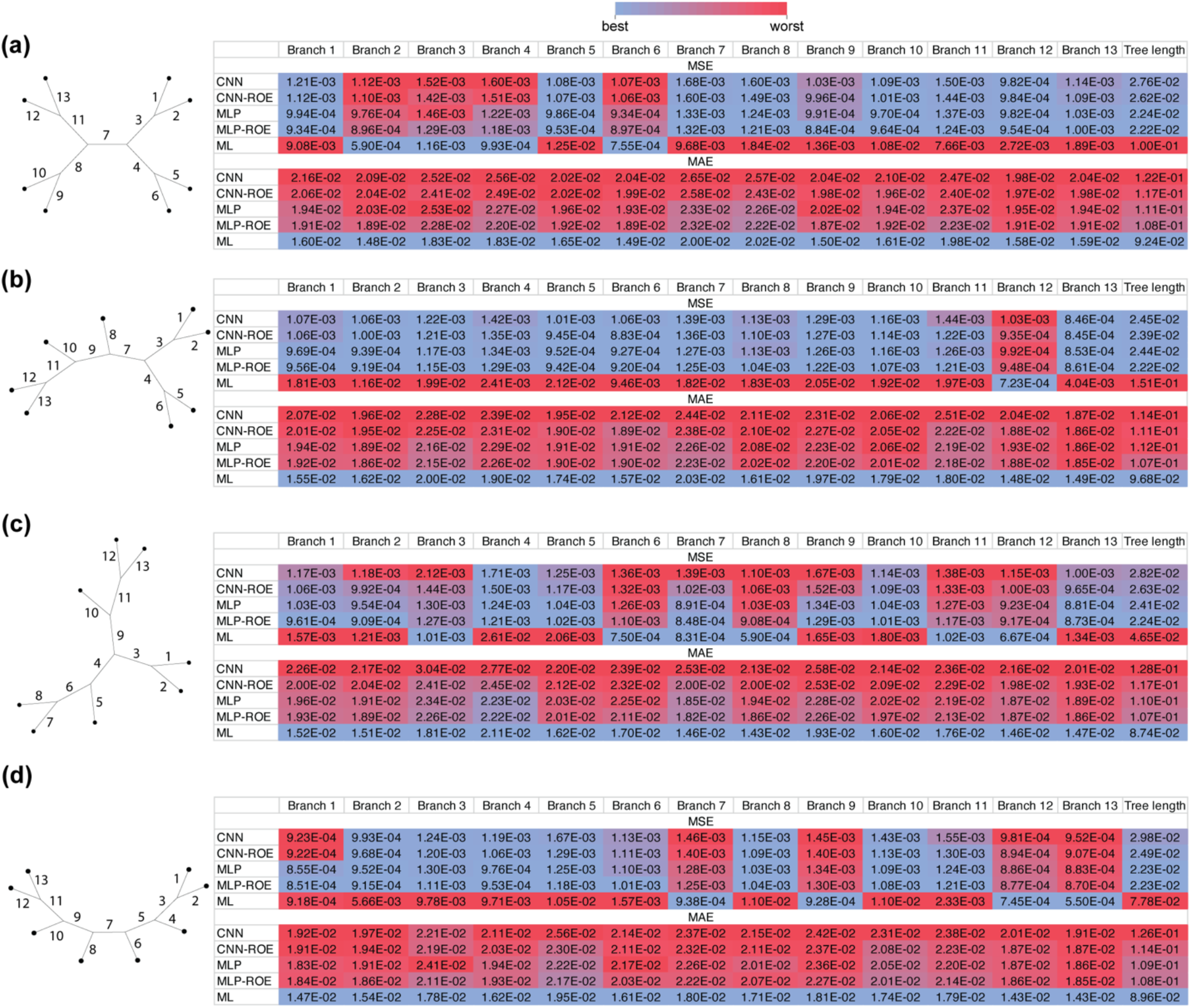
Effects of the shape balance on accuracy of estimated branch length using Mean Squared Error (MSE) and Mean absolute Error (MAE) metrics across methods. (**a**) The table represents a comparison of different methods’ performance for each branch (column names in the tables correspond to the numbered branches of a tree) and total tree length for balanced unrooted tree; panels (**b**) through (**d**) show the same for trees with increasing degrees of imbalance, with the most imbalanced tree, i.e. tree with pectinate (aka caterpillar) shape shown in (**d**). Color scheme ranks MSE or MAE metrics across all methods for a given branch (or total tree length), with the best values for a given branch shown in blue and the worst values shown in red. CNN: convolutional neural network; CNN-ROE: convolutional neural network – regression of observed on estimated values; MLP: multilayer perceptron; MLP-ROE: multilayer perceptron – regression of observed on estimated values; ML: maximum likelihood.

**Figure 6:**
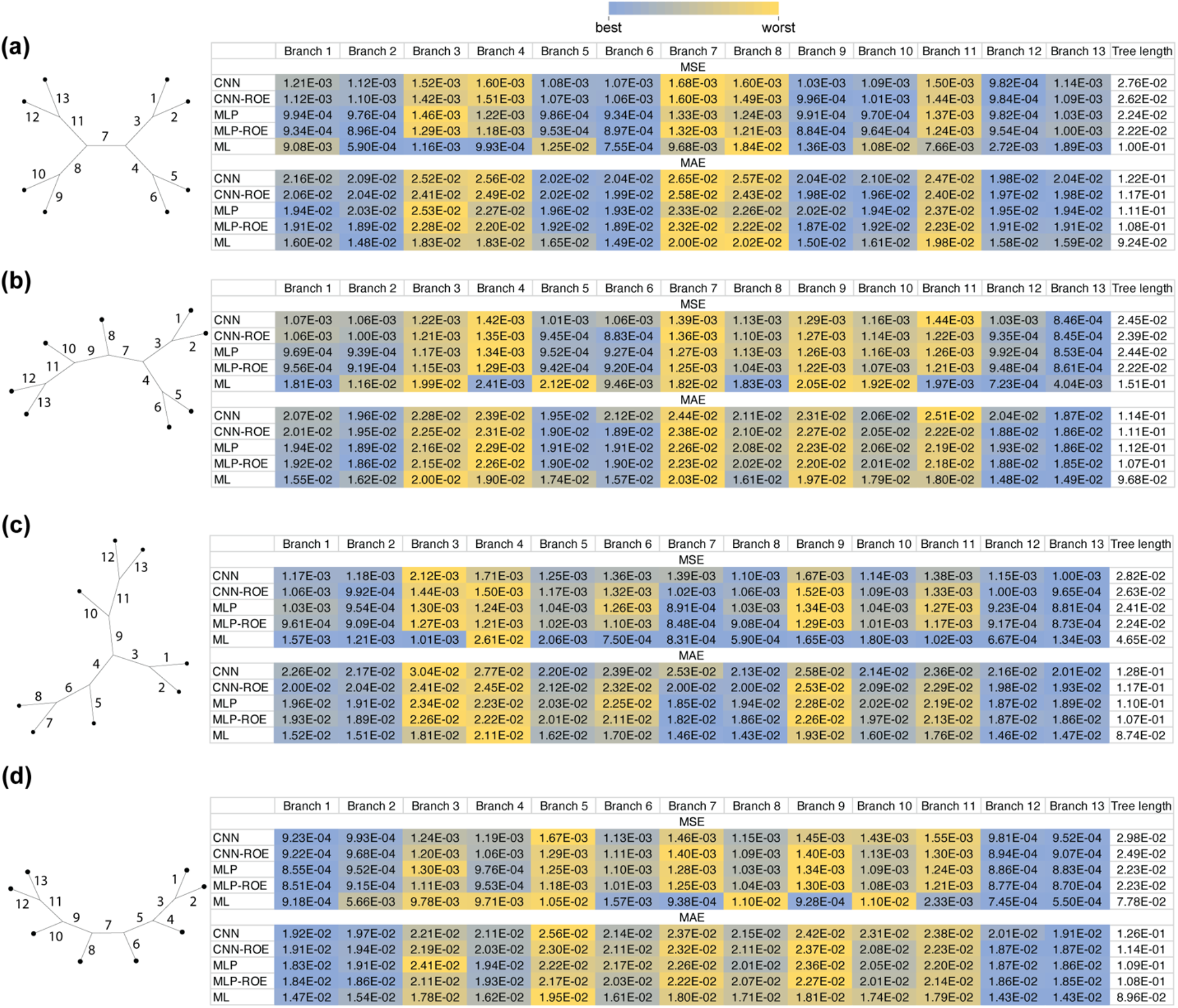
Effects of the shape balance on accuracy of estimated branch length using Mean Squared Error (MSE) and Mean absolute Error (MAE) highlighting differences between branches. Same is **Fig. 5**, but with the color scheme showing which branches receive the lowest (blue) and highest (yellow) error metrics for a given method.

### Concluding remarks and path forward

Here we investigated the possibility of deep learning models to estimate branch lengths on a fixed tree topology. Our ANNs showed excellent performance under a range of simulation conditions. Although ML is a highly accurate method for estimating branch lengths (Schwartz and Mueller 2010), in the majority of conditions considered here, including those regions of parameter space that may cause bias, ANNs provided superior branch length estimates than ML. Additionally, in this study, we did not conduct any comparisons of our ANNs with Bayesian methods, as the latter has been demonstrated to be less reliable than ML for estimating branch lengths (Schwartz and Mueller 2010), as performance can be highly dependent on the choice of branch length prior distribution (Nelson et al. 2015). Due to the flexibility of artificial neural network architectures, such that they can be formulated to simultaneously execute classification and regression tasks, they could potentially be extended to co-estimate tree topology and branch lengths from MSAs. Since, ANNs are accurate estimators individually for both of these fundamental phylogenetic parameters (i.e. topology (Suvorov et al. 2020) and branch lengths as we showed in this study), it is possible that ANNs will exhibit similarly impressive accuracy when these parameters are estimated together, and future studies should explore this possibility. One current limitation of the ANN architectures examined here is that they cannot scale to larger numbers of taxa. However, explorations of alternative ANN architectures, or approaches that combined neural networks with aspects of more traditional tree optimization methods (Azouri et al. 2021), may in due course overcome these limitations.

The encouraging performance of our implementations of ANNs suggest that these and similar methods can be extended to perform fossil calibrations, i.e., inference of branch lengths in absolute time. In principle, it is straightforward to incorporate fossil calibration data into the inputs for an ANN, along with multiple sequence alignments or summaries thereof, to estimate absolute divergence times on a fixed topology, provided that one can generate appropriate training data modeling the process that generates absolute branch length conditioned on the available calibration points. Also, it would be important to investigate the effects of indels on branch length estimation. Our previous study (Suvorov et al. 2020) showed that indel information can be effectively utilized toward more accurate tree topology inference within a machine learning framework. Thus, we expect that ANNs, which can implicitly construct indel models from training MSA by ANNs, may have the potential to improve branch length estimation relative to traditional methods that ignore indel information. Indel events provide useful information for evolutionary inference for highly diverged sequences (Rokas and Holland 2000; Loewenthal et al. 2021) or in scenarios where the number of substitutions is insufficient for reliable estimation (Redelings and Suchard 2007), so indels could have a positive effect on branch length estimation for trees with long or short branches.

## Acknowledgements

We thank Jeff Thorne for valuable feedback and helpful suggestions. This work was funded by the National Institutes of Health under award number R35GM138286.

## Author Contributions

AS and DRS conceptualized the research. AS analyzed the data. AS and DRS wrote the paper.

## Data availability

R and Python scripts can be found at GitHub (https://github.com/antonysuv/tree_branch). Training and test MSA, and branch length datasets as well as trained machine learning models and additional figures for each simulation experiment are deposited on Figshare (https://doi.org/10.6084/m9.figshare.21514272.v2).

