## Supplementary figures and images for "Reliable estimation of tree branch lengths using deep neural networks"

### supplementary fig. S1

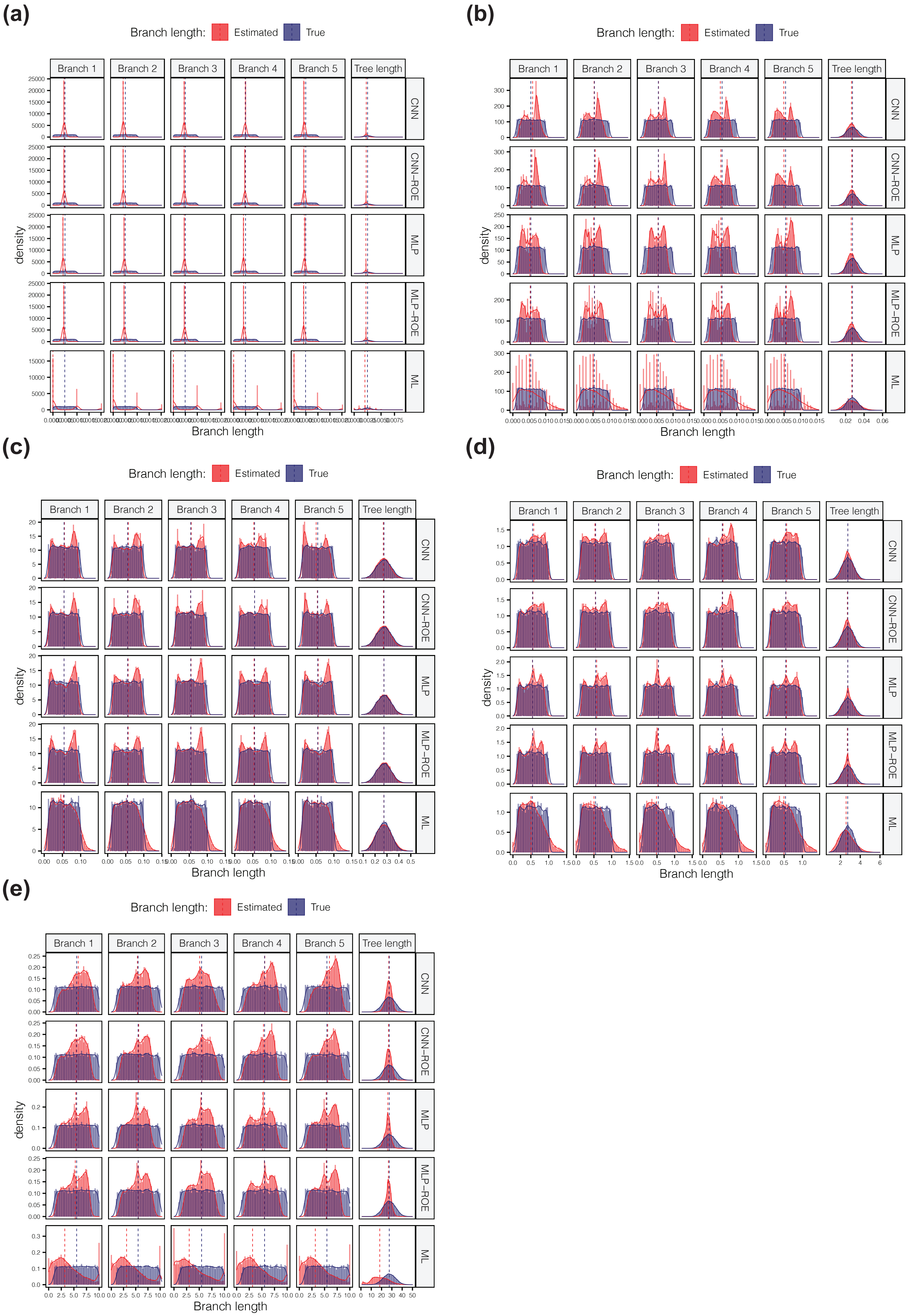

### supplementary fig. S2

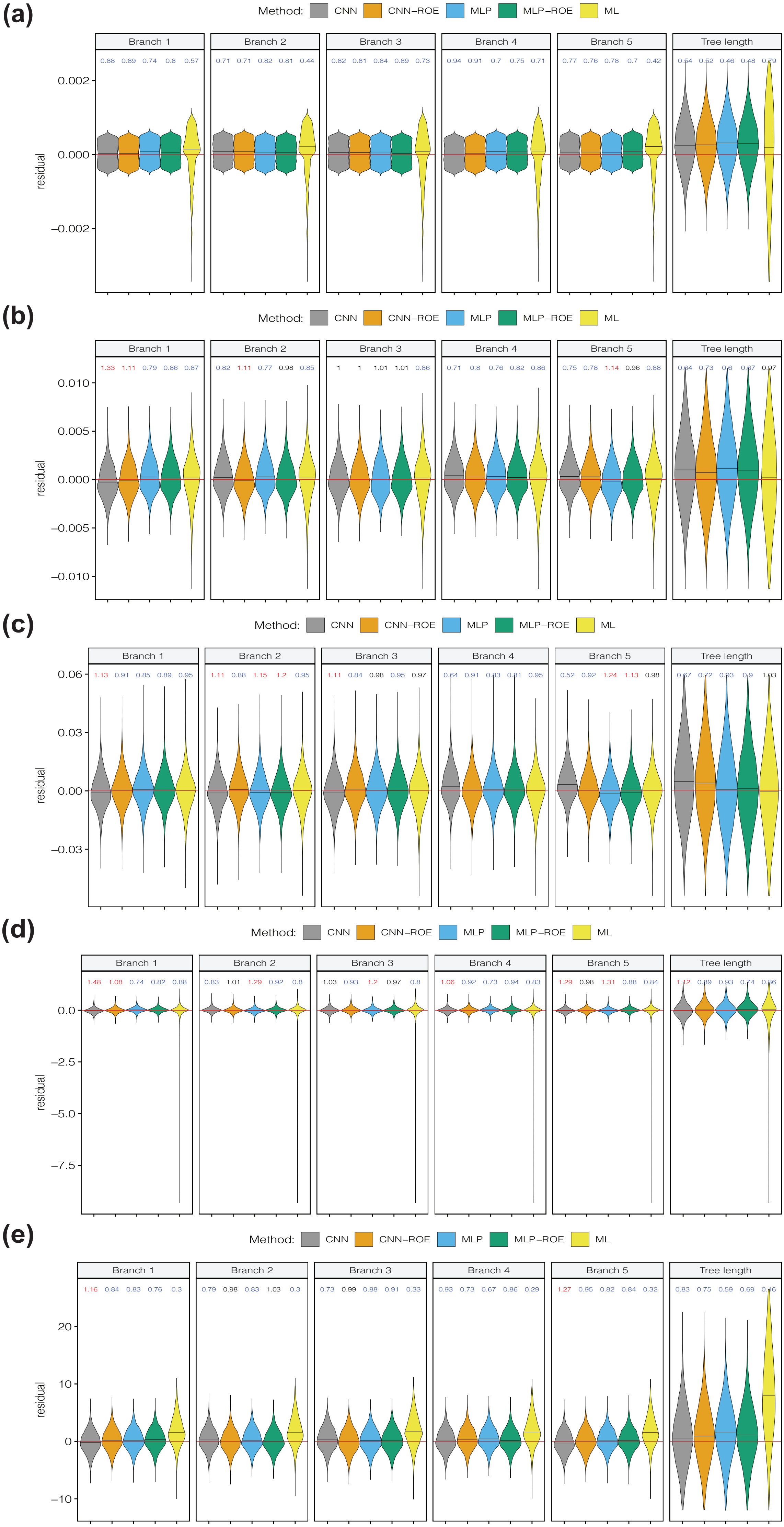

### supplementary fig. S3

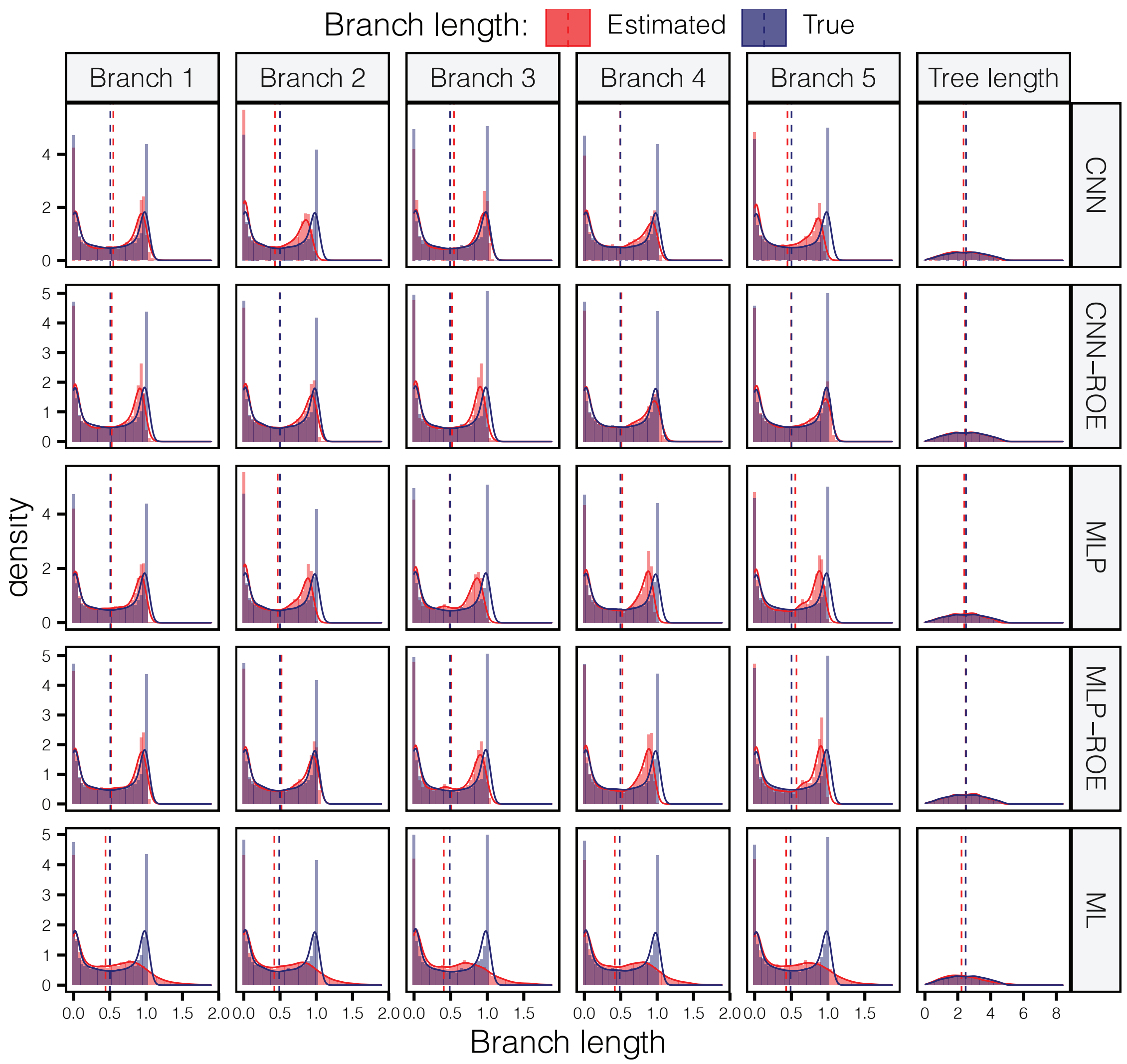

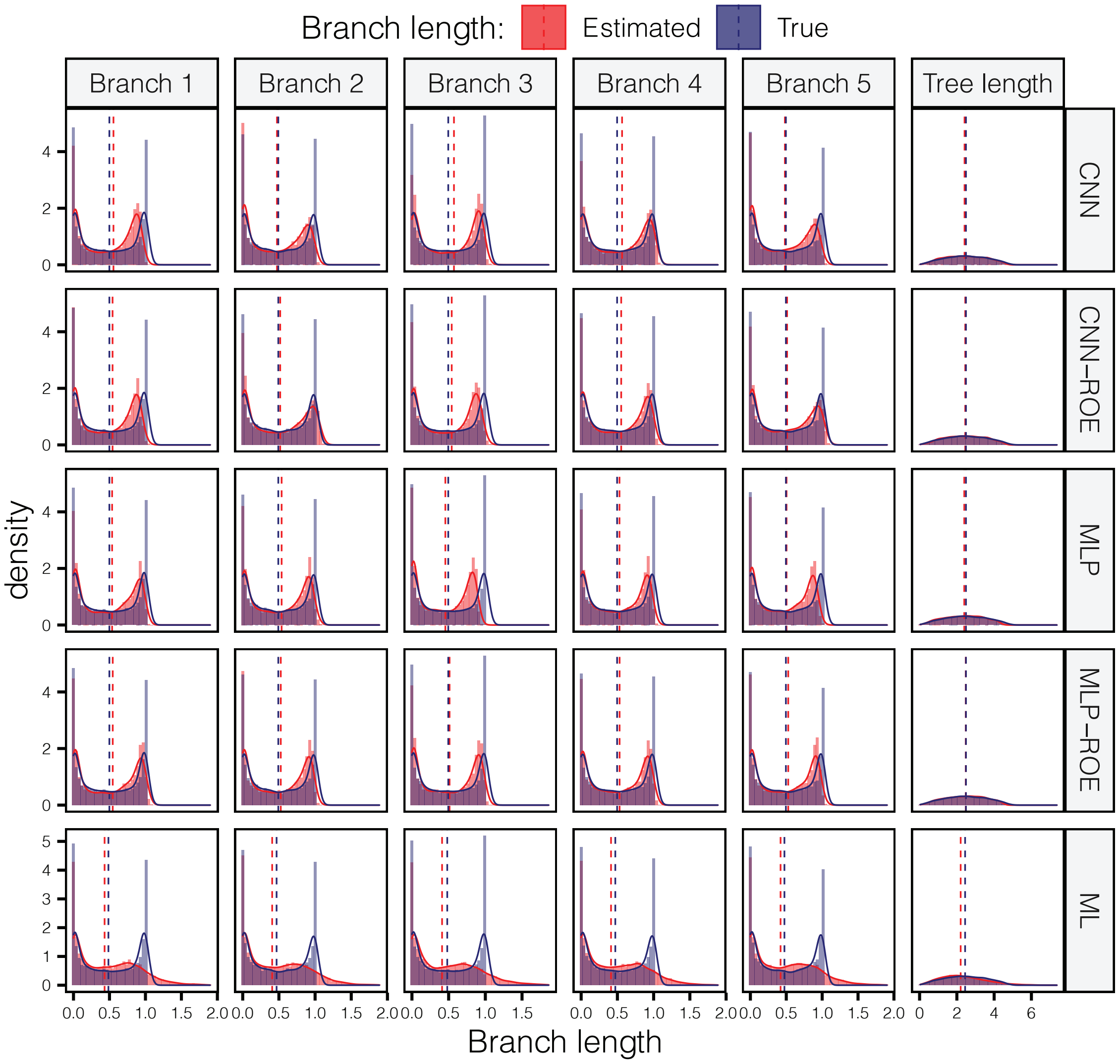

### supplementary fig. S4

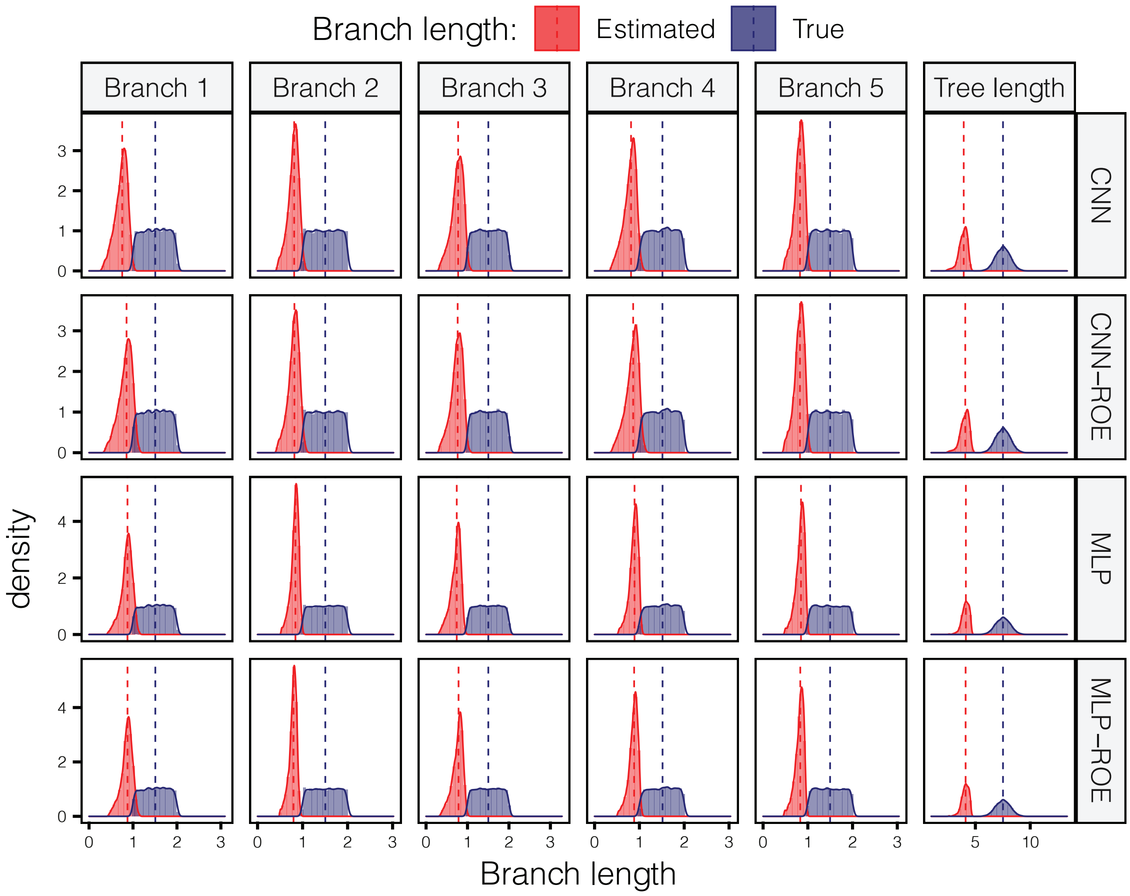

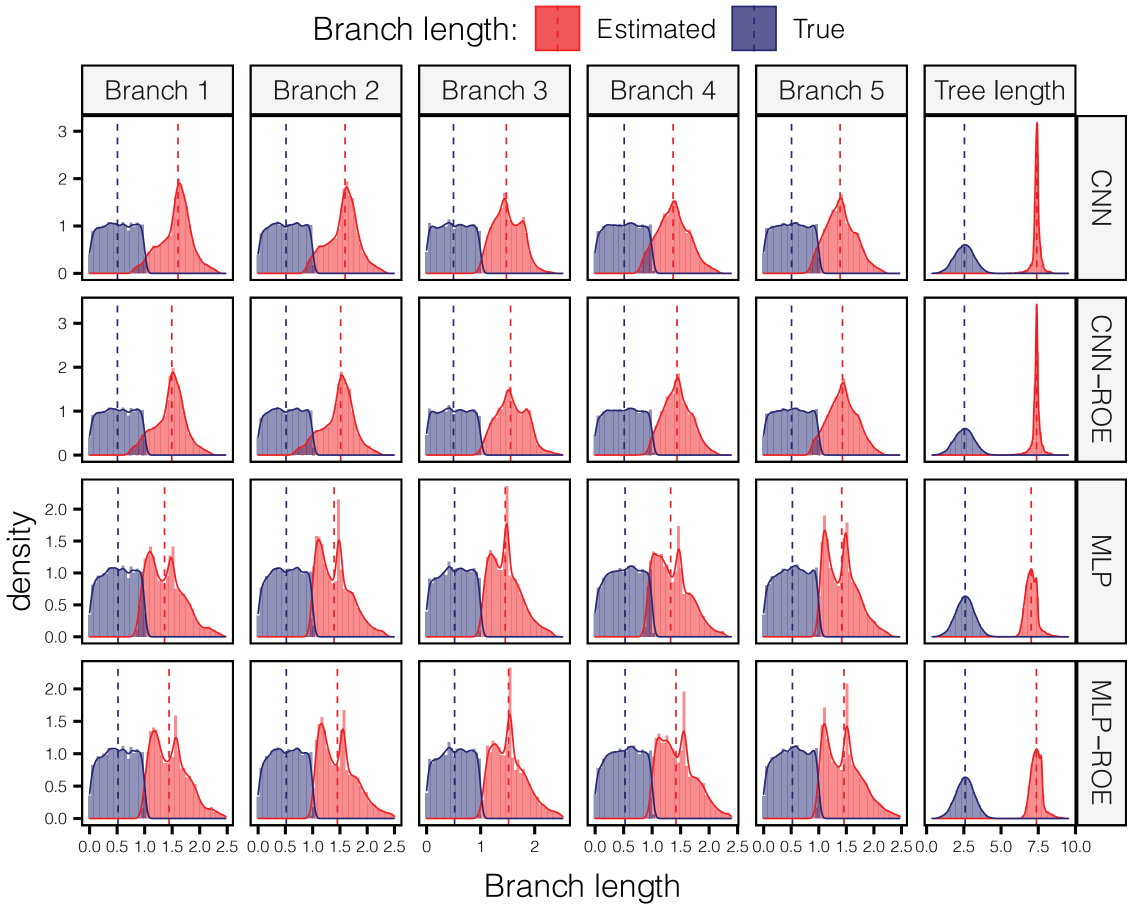
