## supplementary figure and table legends for "Reliable estimation of tree branch lengths using deep neural networks"

**Supplementary Figure Legends**

**Fig. S1: Comparison of predicted and true branch length distributions generated by different uniform distributions.** Branch lengths were generated by: (**a**) $U\left( 0, 0.001 \right)$; (**b**) $U\left( 0.001,0.01 \right)$; (**c**) $U\left( 0.01, 0.1 \right)$; (**d**) $U\left( 0.1, 1 \right)$ ; (**e**) $U\left( 1, 10 \right)$. Density plots represent true (blue) and predicted (red) branch length distributions. Dashed lines mark the position of the median. CNN = convolutional neural network; CNN-ROE = convolutional neural network – regression of observed on estimated values; MLP = multilayer perceptron; MLP-ROE = multilayer perceptron – regression of observed on estimated values; ML = maximum likelihood.

**Fig. S2: Comparison of residual distributions for branch length distributions generated by different uniform distributions.** Branch lengths were generated by: (**a**) $U\left( 0, 0.001 \right)$; (**b**) $U\left( 0.001,0.01 \right)$; (**c**) $U\left( 0.01, 0.1 \right)$; (**d**) $U\left( 0.1, 1 \right)$ ; (**e**) $U\left( 1, 10 \right)$. The violin plot that shows the distribution of residuals (i.e. difference between true and inferred branch lengths) for each method. The value above each violin represents the ratio of overestimated and underestimated branches. The values of ~1 indicate an equal number of over- and underestimates, <1 indicate the underestimation is more common, and >1 indicates that overestimation is more common. The colored values represent statistically significant underestimation (blue) or overestimation (red). The horizontal red line marks 0. The black horizontal line within each violin shows the median.

**Fig. S3: Comparison of predicted and true branch length distributions generated within branch length heterogeneity space (BL-space).** The results were obtained under (**a**) JC and (**b**) GTR substitution models. Density plots represent true (blue) and predicted (red) branch length distributions. Dashed lines mark the position of the median. CNN = convolutional neural network; CNN-ROE = convolutional neural network – regression of observed on estimated values; MLP = multilayer perceptron; MLP-ROE = multilayer perceptron – regression of observed on estimated values; ML = maximum likelihood.

**Fig. S4: Misspecification of the distribution of branch lengths used during training.** (**a**) ANNs were trained on MSAs simulated using trees with branch lengths sampled from $U\left( 0.1, 1 \right)$ and tested on $U\left( 1, 2 \right)$. (**b**) ANNs were trained on MSAs simulated using trees with branch lengths sampled from $U\left( 1, 2 \right)$ and tested on $U\left( 0.1, 1 \right)$. Density plots represent true (blue) and predicted (red) branch length distributions. Dashed lines mark the position of the median. CNN = convolutional neural network; CNN-ROE = convolutional neural network – regression of observed on estimated values; MLP = multilayer perceptron; MLP-ROE = multilayer perceptron – regression of observed on estimated values.

**Supplementary Table Legends**

**Table S1: Results from “Uniform” experiments (see Table 1).** The name of each experiment follows this convention: *experiment_unif_{min}_{max}_{number of taxa}_{substitution model}*. The “*min*” and “*max*” correspond to the minimum and maximum parameter values of a uniform distribution. The following values were computed for each branch and tree length: mean squared error (MSE); mean absolute error (MAE); Spearman’s correlation coefficient (rho); Spearman’s correlation test *P* value (P_rho); bias (Bias); Kolmogorov-Smirnov test statistic *D* (D); Kolmogorov-Smirnov test *P* value (P_D). CNN = convolutional neural network; CNN-ROE = convolutional neural network – regression of observed on estimated values; MLP = multilayer perceptron; MLP-ROE = multilayer perceptron – regression of observed on estimated values; ML = maximum likelihood.

**Table S2: Results from “Exponential” experiments (see Table 1).** The name of each experiment follows this convention: *experiment_exp_{rate}_{number of taxa}_{substitution model}*. The “*rate*” corresponds to the rate parameter of a uniform distribution. The following values were computed for each branch and tree length: mean squared error (MSE); mean absolute error (MAE); Spearman’s correlation coefficient (rho); Spearman’s correlation test *P* value (P_rho); bias (Bias); Kolmogorov-Smirnov test statistic *D* (D); Kolmogorov-Smirnov test *P* value (P_D). CNN = convolutional neural network; CNN-ROE = convolutional neural network – regression of observed on estimated values; MLP = multilayer perceptron; MLP-ROE = multilayer perceptron – regression of observed on estimated values; ML = maximum likelihood.

**Table S3: Results from “Branch-length heterogeneity space (BL-space)” experiments (see Table 1).** The name of each experiment follows this convention: *experiment_mixb_{number of taxa}_{substitution model}*. The “*mixb*” denotes mixture beta distribution. The following values were computed for each branch and tree length: mean squared error (MSE); mean absolute error (MAE); Spearman’s correlation coefficient (rho); Spearman’s correlation test *P* value (P_rho); bias (Bias); Kolmogorov-Smirnov test statistic *D* (D); Kolmogorov-Smirnov test *P* value (P_D). CNN = convolutional neural network; CNN-ROE = convolutional neural network – regression of observed on estimated values; MLP = multilayer perceptron; MLP-ROE = multilayer perceptron – regression of observed on estimated values; ML = maximum likelihood.

**Table S4: Results from “Birth-death” experiments (see Table 1).** The name of each experiment follows this convention: *experiment_bd_{relative extinction}_{age of the root node}_{clock rate}_{substitution model}*. The “*bd*” denotes birth-death model. The following values were computed for each branch and tree length: mean squared error (MSE); mean absolute error (MAE); Spearman’s correlation coefficient (rho); Spearman’s correlation test *P* value (P_rho); bias (Bias); Kolmogorov-Smirnov test statistic *D* (D); Kolmogorov-Smirnov test *P* value (P_D). CNN = convolutional neural network; CNN-ROE = convolutional neural network – regression of observed on estimated values; MLP = multilayer perceptron; MLP-ROE = multilayer perceptron – regression of observed on estimated values; ML = maximum likelihood.

**Table S5: Results from “Model misspecification” experiments (see Table 1).** The name of each experiment follows this convention: *experiment_exp_{rate}_{number of taxa}_{substitution model of testing dataset/substitution model of training dataset}*. The “*rate*” corresponds to the rate parameter of a uniform distribution. The following values were computed for each branch and tree length: mean squared error (MSE); mean absolute error (MAE); Spearman’s correlation coefficient (rho); Spearman’s correlation test *P* value (P_rho); bias (Bias); Kolmogorov-Smirnov test statistic *D* (D); Kolmogorov-Smirnov test *P* value (P_D). CNN = convolutional neural network; CNN-ROE = convolutional neural network – regression of observed on estimated values; MLP = multilayer perceptron; MLP-ROE = multilayer perceptron – regression of observed on estimated values; ML = maximum likelihood.

**Table S6: Results from “Misspecified branch length distribution” experiments (see Table 1).** The name of each experiment follows this convention: *experiment_unif_{number of taxa}_{substitution model}_train_{min}_{max}_test_{min}_{max}*. The “*min*” and “*max*” correspond to the minimum and maximum parameter values of uniform distributions that were used to simulate training and testing datasets. The following values were computed for each branch and tree length: mean squared error (MSE); mean absolute error (MAE); Spearman’s correlation coefficient (rho); Spearman’s correlation test *P* value (P_rho); bias (Bias); Kolmogorov-Smirnov test statistic *D* (D); Kolmogorov-Smirnov test *P* value (P_D). CNN = convolutional neural network; CNN-ROE = convolutional neural network – regression of observed on estimated values; MLP = multilayer perceptron; MLP-ROE = multilayer perceptron – regression of observed on estimated values; ML = maximum likelihood.

**Table S7: Results from “Exponential” experiments (see Table 1).** The name of each experiment follows this convention: *experiment_exp_{rate}_{number of taxa}_{substitution model}_{balance} {tree topology in newick}*. The “*rate*” corresponds to the rate parameter of a uniform distribution. The value of “*balance*” indicates tree topology imbalance, where a bigger number corresponds to larger imbalance. The following values were computed for each branch and tree length: mean squared error (MSE); mean absolute error (MAE); Spearman’s correlation coefficient (rho); Spearman’s correlation test *P* value (P_rho); bias (Bias); Kolmogorov-Smirnov test statistic *D* (D); Kolmogorov-Smirnov test *P* value (P_D). CNN = convolutional neural network; CNN-ROE = convolutional neural network – regression of observed on estimated values; MLP = multilayer perceptron; MLP-ROE = multilayer perceptron – regression of observed on estimated values; ML = maximum likelihood.

**Table S8: Variance of mean squared errors (MSE) and mean absolute errors (MAE) across all branches in 8-taxon trees.** The value of balance indicates tree topology imbalance, where a bigger number corresponds to larger tree imbalance. CNN = convolutional neural network; CNN-ROE = convolutional neural network – regression of observed on estimated values; MLP = multilayer perceptron; MLP-ROE = multilayer perceptron – regression of observed on estimated values; ML = maximum likelihood.
